# Quantifying the dynamics of pig movements improves targeted disease surveillance and control plans

**DOI:** 10.1101/2019.12.29.886408

**Authors:** Gustavo Machado, Jason Ardila Galvis, Francisco Paulo Nunes Lopes, Joana Voges, Antônio Augusto Rosa Medeiros, Nicolas Cespedes Cárdenas

## Abstract

Tracking animal movements over time can fundamentally determine the success of disease control interventions throughout targeting farms that are tightly connected. In commercial pig production, animals are transported between farms based on growth stages, thus it generates time-varying contact networks that will influence the dynamics of disease spread. Still, risk-based surveillance strategies are mostly based on a static network. In this study, we reconstructed the static and temporal pig networks of one Brazilian state from 2017 to 2018, comprising 351,519 movements and 48 million transported pigs. The static networks failed to capture time-respecting movement pathways. Therefore, we propose a time-dependent network susceptible-infected (SI) model to simulate the temporal spread of an epidemic over the pig network globally through the temporal movement of animals among farms, and locally with a stochastic compartmental model in each farm, configured to calculate the minimum number of target farms needed to achieve effective disease control. In addition, we propagated disease on the pig temporal network to calculate the cumulative contacts as a proxy of epidemic sizes and evaluated the impact of network-based disease control strategies. The results show that targeting the first 1,000 farms ranked by degree would be sufficient and feasible to diminish disease spread considerably. Our finding also suggested that assuming a worst-case scenario in which every movement transmit disease, pursuing farms by degree would limit the transmission to up to 29 farms over the two years period, which is lower than the number of infected farms for random surveillance, with epidemic sizes of 2,593 farms. The top 1,000 farms could benefit from enhanced biosecurity plans and improved surveillance, which constitute important next steps in strategizing targeted disease control interventions. Overall, the proposed modeling framework provides a parsimonious solution for targeted disease surveillance when temporal movement data is available.

## Introduction

Between-farm animal movement has historically been associated with disease spread (Ortiz-Pelaez et al., 2006; R. P. Smith et al., 2013). Animal movement data has allowed for the initial development of network-based approaches, which are dominated by the use of static network metrics to identify vulnerable farms. In a contact network, each farm (node) can play a role in the dynamics of disease spread, and some can be classified as vulnerable because they periodically receive infected animals (a high “in” degree); therefore, the identification of these farms is more important in scenarios of the absence or low prevalence of disease (Qi et al., 2019). Such locations may serve as sentinels. On the other hand, farms that are more likely to spread disease throughout their contact network could be targeted, by which restricting or monitoring outgoing movements could effectively reduce disease spread (Cárdenas et al., 2018; Payen et al., 2019). Ultimately, the use of network-based metrics to select farms for isolation or surveillance schemes has shown great value (Mohr et al., 2018; Firestone et al., 2019; Payen et al., 2019). This is a direct consequence of the availability of animal movement data (Büttner et al., 2016; Firestone et al., 2019); however, methodologies have not yet evolved as expected. Most published approaches make use of static networks to describe animal movement and subsequently make inferences about outbreaks and epidemics (Büttner et al., 2013, 2018; Marquetoux et al., 2016; Motta et al., 2017; Alarcón et al., 2019; Kinsley et al., 2019). However, the temporal anomalies and seasonality of movements can only be identified if movement paths are aggregated to smaller time windows (Cárdenas et al., 2018; Augusta et al., 2019; Sterchi et al., 2019), which may not be sufficient to make inferences about disease dynamics within the contact networks. Disease transmission is not restricted to the historical record of movement (Firestone et al., 2019; Qi et al., 2019; Sterchi et al., 2019); namely, when disease transmission actually occurs by the movement of infected animals is likely to be an event in time that is dependent on when animals are sent and, more importantly, when they are received at the final destination, and this rationale has been largely overlooked (Chaters et al., 2019; Sterchi et al., 2019). Nevertheless, static networks have been used to measure the effect of targeted farm removal, on the network fragmentation by monitoring giant strongly or weakly connected components, which would create small islands restricting the overall connectivity among farms (Motta et al., 2017; Mekonnen et al., 2019). Other studies have proposed the use of the basic reproduction number (*R*_0_) and have measured the contribution of node removal in disease transmission (Marquetoux et al., 2016; Alarcón et al., 2019; Passafaro et al., 2019), still based on static views of full networks. Therefore, it is recognized that the current neglected temporality of real-world animal movements has been a constant roadblock to the advancement of network epidemiology, especially in the development of methodologies (Pellis et al., 2015).

Stochastic simulations have been used extensively to measure the impact of disease control strategies in future disease spread scenarios (Smith et al., 2017; Funk et al., 2018; Kim et al., 2019; Lanzas et al., 2020). Mathematical models have also been used to experimentally spread disease over contact networks and reveal important dynamic characteristics of epidemics (Miller, 2017; P. Kim and Lee, 2018; Darbon et al., 2019). For example, compartmental models can be used to describe the epidemic spreading in temporal contact networks, which provides an opportunity for tracing outbreaks and contact among nodes over time (C. Guinat et al., 2016; Colman et al., 2019; Ferdousi et al., 2019; Sterchi et al., 2019). Here, we used two years of pig movement data from one Brazilian state to estimate the optimal number of farms for target surveillance by tracking between-farm contact pathways and inferring epidemic sizes throughout disease propagation. We first described the complete pig movements and then challenged the use of static networks against dynamic networks. We proposed a stochastic compartmental model to simulate disease spread in a two-year empirical network while accounting for the temporal order of the movements. Our model formulation allowed the calculation of the minimum number of farms needed to reduce disease spread and approximate the impact of network-based disease control strategies (e.g., farm isolation, movement restriction, enhanced biosecurity) on the expected epidemic sizes. Finally, we applied the proposed model to the time-varying pig network, ultimately to provide a practical background for the identification of targeted farms, which were then mapped and described in regard to their biosecurity and infrastructure.

## Material and methods

### Data collection and entry

The record of pig shipments among all registered farms of one Brazilian state was used to reconstruct contact networks from 2017 until 2018 (SEAPI-RS, 2018). Information about each swine farm included farm identification, production types (breeding, certified-swine-breeder, sow, nursery, wean-to-finisher, finisher, subsistence and others), farms infrastructure (e.g., presence of a fence, presence of a cleaning and disinfection station, see table 2 for the full list), geolocation and farms population capacity. Each pig shipment included date(s), farm of origin and destination, number of animals transported and purpose of movement (weaning, finishing, slaughter or other). Movement data with missing information of farm location, production type, farm of origin or destination were excluded prior to the analysis. Additionally, farms declared inactive (no pigs raised in the past two years or out of business) were not considered in the analysis. Movements from or to other states were also excluded from any analysis. Based in information reported to the local authorities in June 2018 all farms were classified as i) “commercial” farm with active contracts with integrated swine companies, ii) “independent” farm that declared to be commercial but did not have active contract with integrated swine companies and small farm holders that, and iii) “not reported” farms that failed to report to local authorities their pig operation type. The identification of 1,911 (21.4%) not reported farms, resulted in regional policy in which the state authorities issued a request to all farms report pig operation status, within one year, farms that fail to report will have all pig movements blocked until information is shared. Finally, from the total number of active pig farms 11,849, a subset of farms 9,500 had registered geographic coordinates which included commercial, independent and not reported business type, therefore were used for the simulation modeling subsections.

**Table 1.** Description of network terminology and definition applied to pig movements.

| Parameter | Definition | References |
| --- | --- | --- |
| Nodes | A pig farms. | (Wasserman and Faust, 1994) |
| Edge | A link between two nodes. | Wasserman and Faust (1994) |
| Degree (k) | The number of unique contacts to and from a farm. When direction was taken into account, the ingoing and outgoing contacts were separated: the “out” degree is the number of contacts originating from a specific farm, and the “in” degree is the number of contacts directed toward a specific premise. | (Wasserman and Faust, 1994) |
| PageRank | Measure of the Google PageRank, a link analysis algorithm that produces a ranking importance of all the farms from 0 to 1. | (Brin and Page, 1998) |
| Closeness centrality | Closeness centrality measures how many steps are required to access every other vertex from a given node. This measure can be directed as incoming steps or outgoing steps. | (Freeman, 1978) |
| Betweenness | The frequency at which a node is located on the shortest path between any two pairs of nodes in the network. | (Freeman, 1978) |
| Giant weakly connected component (GWCC) | The proportion of nodes that are connected in the largest component when directionality of movement is ignored. | Wasserman and Faust (1994) |
| Giant strongly connected component (GSCC) | The proportion of nodes that are connected in the largest component when directionality of movement is considered. | (Wasserman and Faust, 1994) |

**Table 2.** Description of biosecurity and infrastructure of the targeted farms.

| **Variable** | **Overall proportion** |
| --- | --- |
| Presence of isolation barn(s) | No=77.6%<br>Yes=22.4% |
| Fence around the farm | No=54.7%<br>Yes=45.3% |
| Cleaning and disinfection station(s) | No=72.7%<br>Yes=27.3% |
| Vegetation barrier around the farm | No=86.7%<br>Yes=13.3% |
| Single entry point | No=72.4%<br>Yes=27.6% |
| Bird nets | No=59.6%<br>Yes=40.4% |
| Pig loading and unloading outside the farm's designated clean area | No=61.3%<br>Yes=38.7% |
| Special clothing dedicated for farm use (shower in) | No=77.7%<br>Yes=22.3% |
| Feed bins inside the designated clean area while loading is external | No=60.3%<br>Yes=39.7% |
| ***Vulnerability score \**** |  |
| Low | 0.21% |
| Medium | 47.4% |
| High | 17.8% |
| Not informed | 34.8% |
| ***Veterinary care*** |  |
| From commercial system | 63.5% |
| Independent veterinary | 4.5% |
| Absent | 1.1% |
| Not informed | 30.9% |
| ***Farm type*** |  |
| Nursery | 32.23% |
| Breeding farm | 20.2% |
| Finisher farm | 15.1% |
| Certified swine breeder farm | 5.65% |
| Wean-to-finisher | 1.49% |
| Not informed | 25.33% |
| ***Water sources*** |  |
| Pond | 42.8% |
| Natural water fountain | 22.2% |
| Public source | 9.9% |
| Lake | 1.5% |
| Water cistern | 0.3% |
| Not informed | 23.3% |
| ***Feed source*** |  |
| From production system | 68.7% |
| Own feed mill | 7.8% |
| Not informed | 23.5% |
| ***Destination of dead animals*** |  |
| Compost | 76.0% |
| Incinerator | 0.23% |
| Bury | 0.23% |
| Septic pit | 0.24% |
| Not informed | 23.3% |
| ***Manure destination*** |  |
| Lagoon | 28.9% |
| Septic pit | 2.48% |
| Not informed | 68.62% |
| ***Barn infrastructure*** |  |
| Concrete or bricks | 51.9% |
| Wood | 1.29% |
| Mixed of wood and concrete | 23.4% |
| Not informed | 23.5% |

\*This measures the level of vulnerability the introduction of pathogen into certified-swine-breeder-farms, briefly is a combination of risks measurements form distance to farms to presence of biosecurity and infrastructure of the farm.

### Static and monthly networks reconstruction and analysis

Two directed networks were reconstructed. First, a static network with directed “edges” represented by pig shipments between two “nodes”, where nodes corresponded to pig farms. The static network movements from January 2017 to December 2018 were analyzed by the traditional methodology in which movements are aggregated into one snapshot prior to the reconstruction of the networks (Cárdenas et al., 2018). Second, directed networks were constructed based on a temporal resolution of 30 day windows, in which directed monthly-networks were generated. (Lentz et al., 2016). For both complete and monthly networks, centrality metrics were calculated (Table 1) (Cárdenas et al., 2018). As slaughterhouses constitute sinks and we were interested in potential disease spread among living animals, transports to slaughter were excluded from the network analysis, but are kept for descriptive purposes (see figure 2 and supplement table S1).

**Figure 1.**
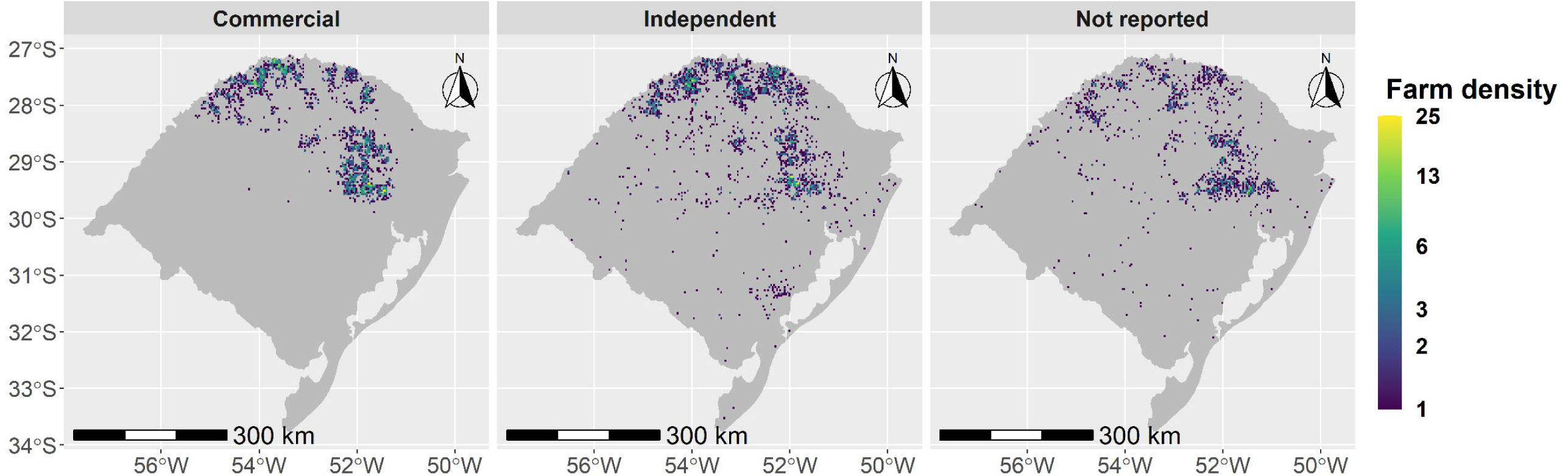
Geographical location of farm density by type of pig operation, commercial, independent, and not reported.

**Figure 2.**
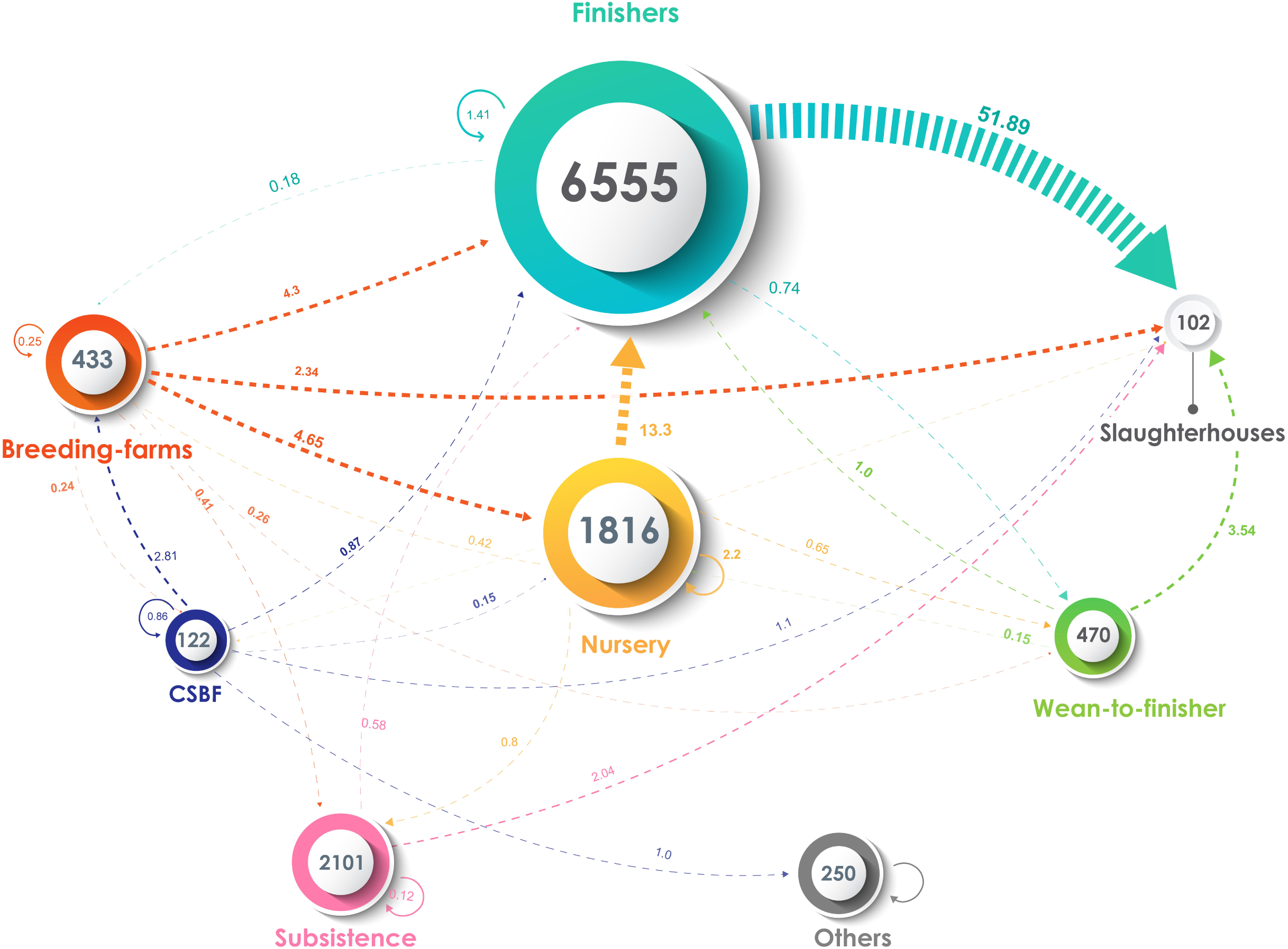
Description of pig movements between all farms types from January 2017 to December 2018. The arrows size shows the percentage of movements and the color is given by the origin of the movements. The size of the circles represented the total number of farms.

### Temporal variability of the network structure

Static networks are considered an approximation of all movement paths for a defined period, e.g., animal movement between pig farms for one month (Büttner et al., 2013; Lentz et al., 2016). This can artificially create accessibility paths between nodes that are not always accessible by all “nodes” in a network, which can lead to an overestimation of the connectivity among the nodes (Holme and Saramäki, 2012; Lentz et al., 2016). To quantify the amounts of such static error, the number of paths in the static view can be compared with the number of paths in the time-series network (Lentz et al., 2013, 2016). This ratio is known as the causal fidelity “c”, where:

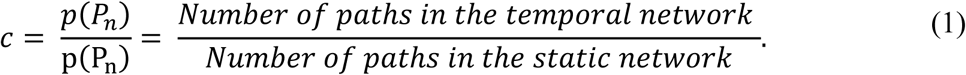

The number of paths is the number of nonzero elements in *p* or *P*. A causal fidelity of one means that the temporal network is well represented by its static network, while networks with lower causal fidelity values would not represent the temporal paths well because most of the paths are not causal. Here, we calculated the causal fidelity on timescales that are typically used to investigate animal movement data: monthly, 6 months and 1 year and for the full two years of data that were used to reconstruct directed networks (Lentz et al., 2016). We also calculated the edge loyalty (*θ*), which measures the fraction of preserved edges (*θ*) of a given premise between two consecutive years, *t* − 1 and *t*. To quantify 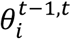, we define 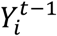 as the set of edges from the premise *i* in the time *t* − 1 and 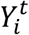 as the set of edges from the premise *i* at time *t*. Then, 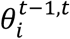 is given by the Jaccard Index in equation 2 (Valdano et al., 2015; Schulz et al., 2017).

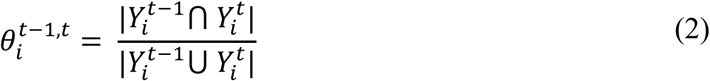

For the node loyalty, we use the same Jaccard Index,

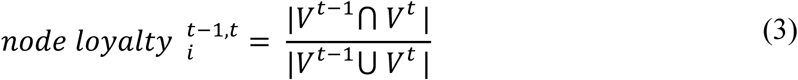

where 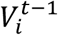 is the set of farms that are active in period *t* − 1, and 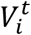 is the set of farms that moved at least one animal at time *t* (Valdano et al., 2015; Schulz et al., 2017).

### The minimum number of farms needed for effective control disease spreading process over time-varying network

We developed a stochastic susceptible-infectious (SI) epidemic model to calculate the minimum number of farms needed to effectively reduce the number of new cases expected to spread through the contact network. The number of infections after the removal of farms raked by network metrics betweenness, closeness, page rank and degree, random removals (similar to the current state of farm surveillance), and without farm removal were the main model outputs. Ultimately, this approximates the impact of network-based target surveillance, which often include farm visits, sample collection, etc., to a feasible number of farms. This model framework integrates infection dynamics in each subpopulation as continuous-time Markov chains using the Gillespie stochastic simulation algorithm (Widgren et al., 2019). Transitions from *S* → *I* was in a two-stage model with a one-way transition depending on 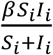 with *β* as the transmission rate.

The completely stochastic simulation can be summarized as follows: the empirical movement data from 2017 until 2018 was used to simulate disease spread over a time-varying network; here, we assume an initial herd prevalence of 10% and 0.1% of infected pigs in each farm, in which disease was allowed to be transmitted between and within farms. For the within-farm transmission rate, we use a set of *β* values of 0.01, 0.05, 0.1 and 0.2, which can also be considered the product of the contact rates and the transmission probability between animals into the farms. Another assumption was that the contact rates among pigs were homogenous. For the between-farm transmission, we considered the observed number of pigs that moved between farms and the population size at each farm. As an initial condition, model simulation started with 950 infected farms (10% prevalence) at time “0”. Here, we proposed two realistic strategies to seed initial infected nodes. First, pig farms were selected at random from all farms with at least one movement from 2017 to 2018. For the second strategy, infected farms were chosen proportionally to the number of farms in commercial, independent, and not reported. This allowed the initial infection to be seeded within all types of pig operations. The comparison of both disease introduction approaches allowed us to test a more realistic scenario since disease can emerge at any location of the study area and in any pig operation (see Figure 1). The simulated control intervention was based only on the sequential removal of farms from the time “0” based on the static network metrics described above. While farms were removed, the number of new cases per 100 susceptible farms was calculated. We chose an arbitrary number of 2,000 herds to be removed, and at the end of each simulation scenario the incidence of new cases *I* was calculated as follows:

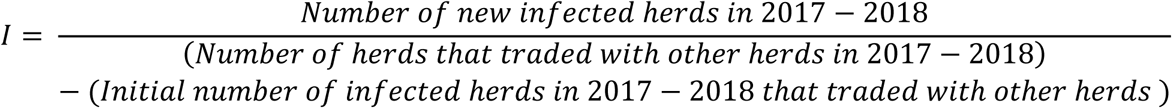

Infection was seeded into all candidate farms randomly 100 times, from which simulations were run and the median incidence was derived for each run. For a sensitivity analysis, we ran simulations with a diverse set of *β* parameters (see supplementary figure S3).

### Cumulative contacts as a proxy of epidemic sizes

The modeling setup described by (Payen et al., 2019) was adapted to identify the network-based metric that can produce the smallest epidemic size distribution at the end of the two years of pig movements, and consequently allow the generation of a farm hot-list to use to target surveillance activities. This approach is based on the classic deterministic SI spreading model that follows movements temporality (Dubé et al., 2008; Nöremark et al., 2011; Konschake et al., 2013). Thereafter, we will refer to this model as “spread cascade”, which is a temporal representation of the cumulative and consecutive contacts among farms, and therefore can be interpreted as the size of an epidemic (Payen et al., 2019). The hypothetical disease spread followed the chronological order of movements over the two years. In short, when there was a directed interaction, including the three dimensions of time *t*, infected farm *i* and susceptible farms *j*, disease has successfully been transmitted. These events were counted in favor of each farm *i* cascade and the triplet *t, i, j* and added to the temporal links of the stream; see (Payen et al., 2019) for more details of the methodology. Overall, we recorded the daily spread cascades for the two years of data, which are represented by weekly averages and ± 95% CI (see supplementary figure S4), and the accumulated spread cascades sizes for each seed node (9,500 farms) in the network at the end of the two years as a proxy for the worst-case scenario in epidemic propagation (Payen et al., 2019). We used the static network to initially rank all pig farms in descending order of degree, betweenness, PageRank, and closeness and then removed the top 10, 250, 500, 750 and 1,000 farms before starting the model simulation while recording the spread cascade sizes. For comparison with usual surveillance practices, we selected the same number of farms randomly. The contribution on containing the epidemic spread by each network-based target surveillance strategy was evaluated by comparison of the final spread cascade distributions at December 2018 using a nonparametric Kruskal-Wallis test followed by Dunn’s post hoc test, where the p-values below 5% were significant. Once the most efficient network-based surveillance strategy was defined, the biosecurity, infrastructure and population of the selected farms were described.

### Software

The software used for statistical analysis and graphics was R statistical software (versions 3.4.1, 5.1.3) (R Core Team) with R Studio editor using the igraph 1.2.4 (Csardi and Nepusz, 2006), tidyverse 1.2.1(Wickham, 2017), SimInf 6.3.0 (Widgren et al., 2019), sf 0.5-3 (Pebesma, 2018), and brazilmaps 0.1.0 (Siqueira, 2019) packages. Spyder 3.3.3 software for Python version 3.7 and Adobe Illustrator CC 2018 software were also used.

## Results

### Description of pig movements

The total number of active pig farms in the complete database included 11,849 farms, 351,519 movements and more than 48 million pigs transported. The majority, 40.4%, were identified as commercial farms, 38.2% of independent pig producers, and 21.4% failed to report pig operation type (see preliminary table S1 to the description of the population in both pig operation classes). In the commercial farms there were 17 pig production systems with the number of participant farms varying from 10 to 465. While we have described movements into the 102 registered slaughterhouses, they were not accounted for in any network analysis thereafter (see Figure 2). Shipments from commercial to slaughterhouse totaled about 34% of the total movements, while 18% of the movements were from independent farms into slaughterhouses and 14% were between commercial farms. Importantly movements from independent pig producers and from farms that did not report type pig operation types into commercial farms were, 3.35% and 1.97%, respectively. This interaction between non-commercial and commercial farms raises concerns about the introduction and spread of known and unknown diseases circulation into highly connected commercial farms (see supplementary table S1). Among all farms with information about production types 6,555 (55.32%) finisher farms, 1,816 (15.33%) nurseries, 470 (3.97%) wean-to-finisher farms, 433 (3.65%) breeding farms, 122 (1.03%) certified swine breeder farm and 250 (2.11%) other farms, which included insemination stations and isolation units, 2,101 (17.73%) as small pig holders (backyard farms or for subsistence).

More than 51% of movements were from finisher farms to slaughterhouses, with a monthly median of 7,657 shipments. The second largest number of movements from nurseries to finisher farms was 13.51%, with a monthly median of 1,992 transfers. Breeding farms made up for 4.65% of the movements to finisher farms and 4.3% of movements to the nurseries, with 673 monthly median movements. Finally, wean-to-finisher farms sent a small but significant amount

of pigs to slaughter (3.5%), while certified swine breeder farms accounted for 2.8% of movements to slaughter (see Figure 2).

### Monthly network analysis

From the two years of movement data, we reconstructed the monthly static networks. The number of farms, movements and animals varied substantially with a steady increase from July 2017 until October of the same year (see Figure 3). The network diameter revealed a positive increasing trend of 33% from April 2018 through December 2018, December 2018, which decreased thereafter; this pattern was different from the monthly values from the previous year. The monthly analysis showed a positive trend of the GSCC in 2017 and a negative trend in 2018, and the GWCC had a more stable pattern with two consecutive peaks in July and October of 2018. In May 2017 GSCC peaked at 40 farms inflated with mostly commercial farms, and in November 2018 the GWCC had almost 600 farms in which most were also commercial (see supplementary Figure S1 and Table S2 for the two years GSCC and GWCC).

**Figure 3.**
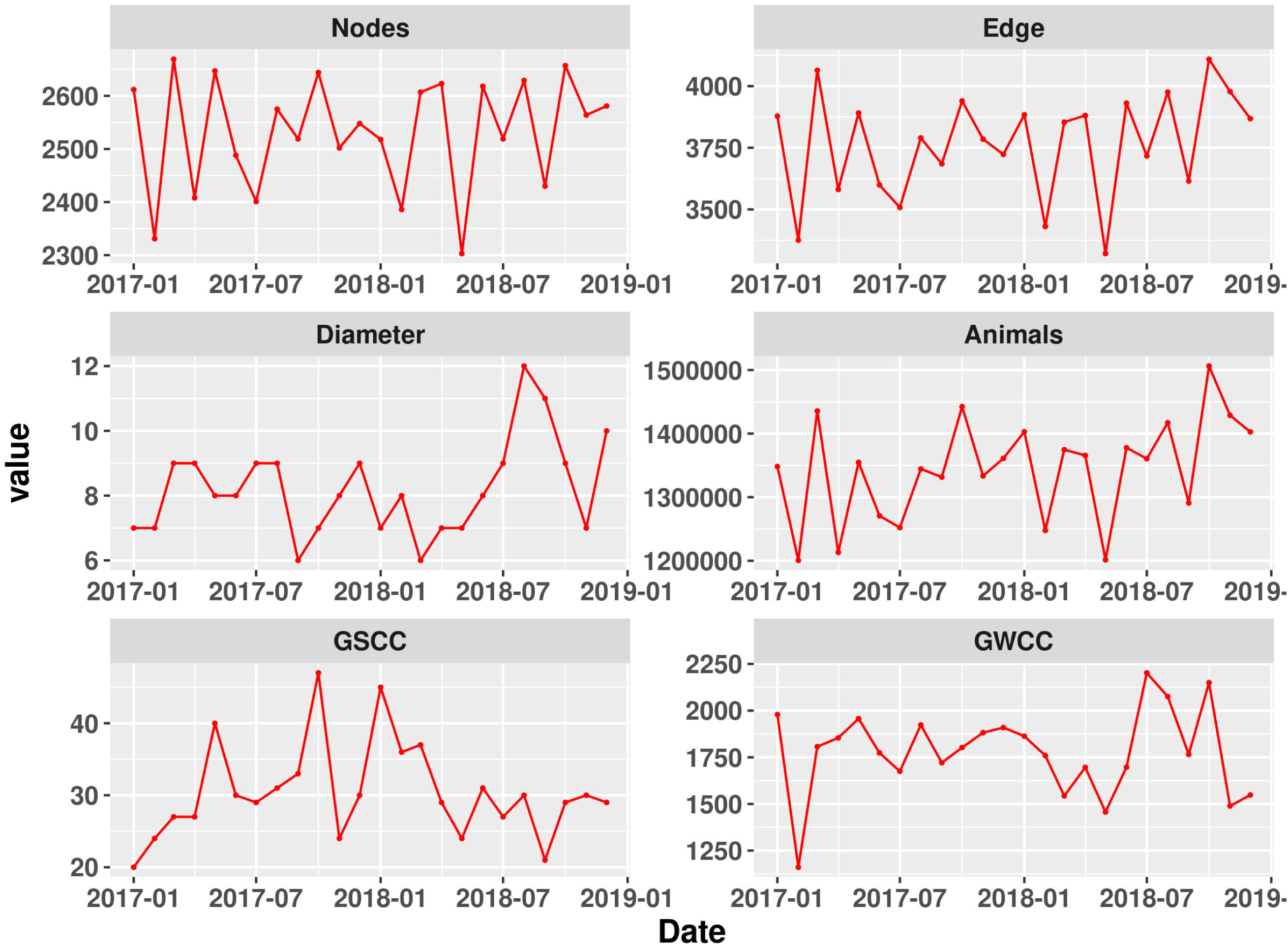
Description of the monthly variation of the network metrics of the pig movements from January 2017 until December 2018.

### Loyalty and causal fidelity

The node loyalty of 2018 compared with 2017 was 0.65, while edge loyalty was 0.19. This means that only 65% of the farms that traded pigs in 2017 were preserved in 2018, and 19% of the edges were preserved between 2017 and 2018. On the other hand, the monthly causal fidelity varied from 0.33 in July 2018 and reached the highest value of 0.55 in July 2017 (see supplementary Figure S2). For the network that evaluated time periods of six months, the causal fidelity was 0.22 for the first half of 2017 and 0.47 for the second half of 2018, and the annual evaluation was in 2017, 0.18 in 2018, and 0.33 for the entire static network. The causal fidelity demonstrated a discrepancy between static and temporal network views. The concept of time does not per se exist in these static networks; thus, the use of traversal network models here would reduce the accuracy toward overestimating disease outbreaks, and likely mislead the selection of farms to undergo network-based surveillance (Lentz et al., 2016). In addition, we calculated the causal error from the causal fidelity, which can be approximated by the inverse of “c” and represents the overall overestimation of disease outbreaks if static networks are used. The first six months of 2017 had the worst results, with overestimations at 4.54 and 1.81 for the July 2018 network.

### Number of farms for targeted surveillance and epidemic sizes

The stochastic runs showed that the removal of 2,000 farms did not completely control the disease from spreading, but it was reduced markedly to a maximum of 1.95 new cases per 100 farms when *β* of 0.2 was used (the worst-case scenario). Since this study was designed to guide decisions in the field, we consulted with the local stakeholders before deciding on the final number of farms to be on the surveillance hot-list (personal communications with Dr. Lopes, Chair of the Animal Movement and Surveillance Department of the state of Rio Grande do Sul). Based on local capacities and the average number of new cases was reduced to as low as 7.88 cases per 100 farms with a *β* of 0.2 when 1,000 farms were removed from the network, the former was set to be the maximum number of feasible farms to be targeted. Furthermore, we compared two infectious seed scenarios: a) infections started at random farms and b) stratified random infections forced to start in proportion to the number of commercial, independent, and not reported. Both have very similar results which reinforced the importance of independent farms in the role of spreading diseases into commercial pig farms and vice versa (Figure 4, also see supplementary figure S3)

**Figure 4.**
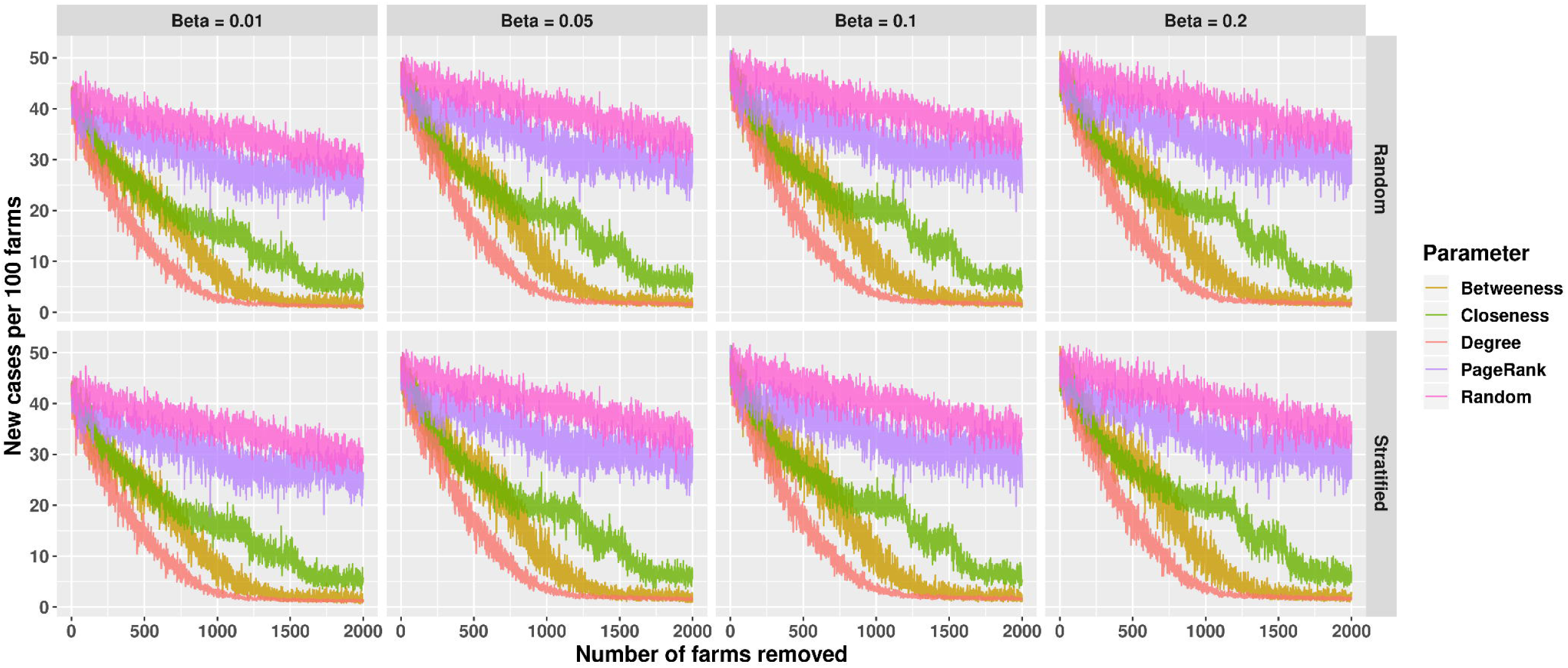
Simulated disease spread dynamics within the swine contact network. Simulations assumed two seed infection scenarios a) random, where 950 pig farms (10% prevalence) were infected at day “0”, and b) stratified, in which the proportion of the 950 pig farms (10% prevalence) was equally distributed by commercial, independent, and not reported farms operation types. The simulations assumed within farm prevalence of 0.1%.

After the removal of the first 1,000 farms with the highest degree, the deterministic model estimated that the maximum outbreak size would involve 29 farms (Figure 5). As expected, random removals resulted in a much larger epidemic size, reaching up to 2,593 farms (Figure 5, also see supplementary table S4 and figure S5).

**Figure 5.**
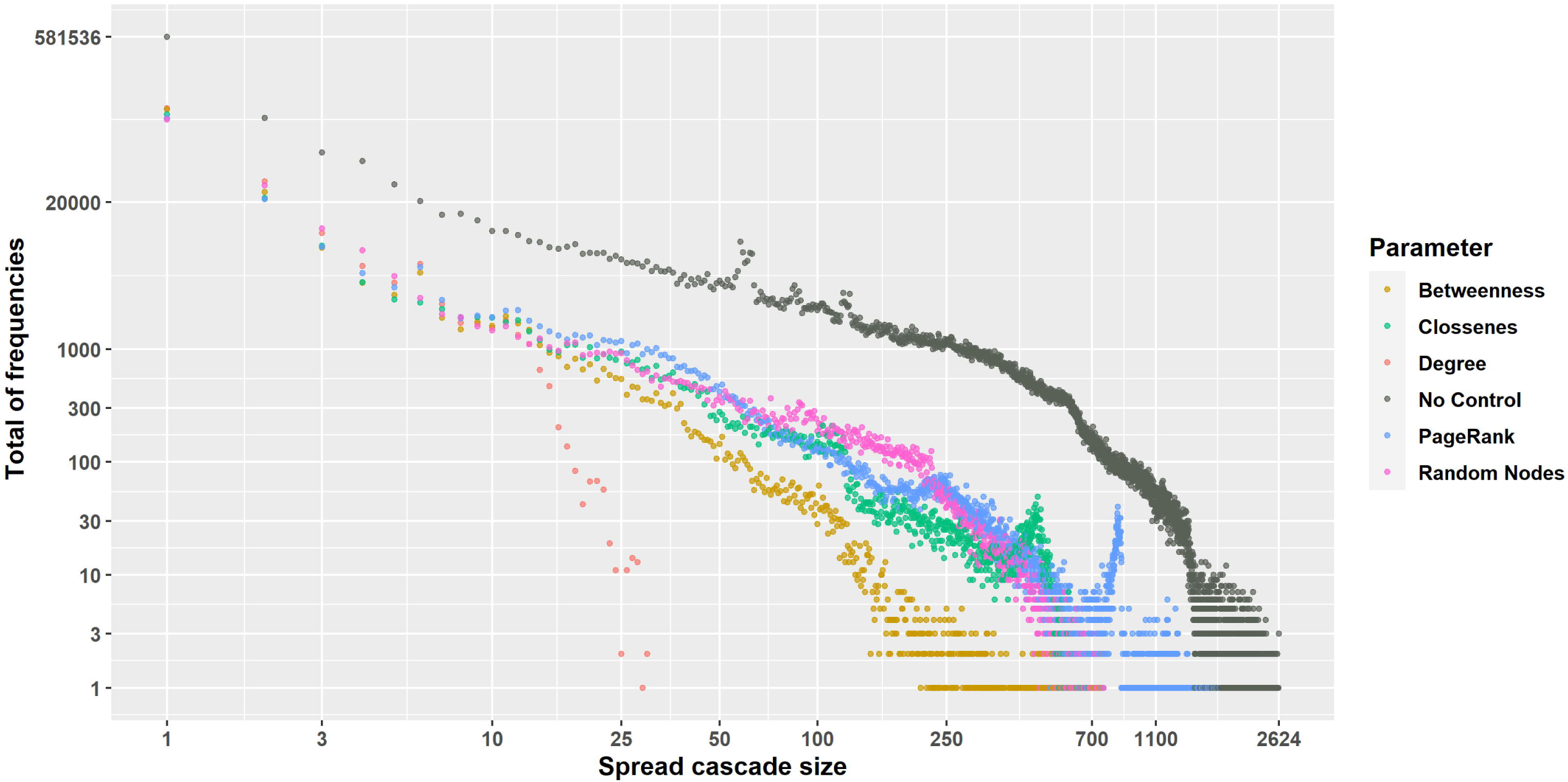
The two years spread cascades sizes for each target-network surveillance strategy. In light blue, the estimated spread cascade, which had a maximum value of 29 farms while not targeting farms, resulted in a cascade of 2,624 farms (black).

### Descriptive analysis of biosecurity, infrastructure and population profile of the targeted farms

Most of the farms within the 1,000 target farms were commercial (55.6%), see Figure 6, with one production system contribution with 138 farms, and 29.7% were independent and 14.7% did not report business operation From the comprehensive list of biosecurity and infrastructure characteristics of the commercial farms, farms had a median of 2 barns (IQR-1-11), capacities for housing were approximately 1,100 pigs (IQR=162-9,925), the median number of piglets was 1,122 (IQR-16-7,229), breeding farms had 4 (IQR=1-18) boars, and the gilts population was 544 (IQR-11-3,044). More details about the farms’ biosecurity and infrastructures can be found in Table 2.

**Figure 6.**
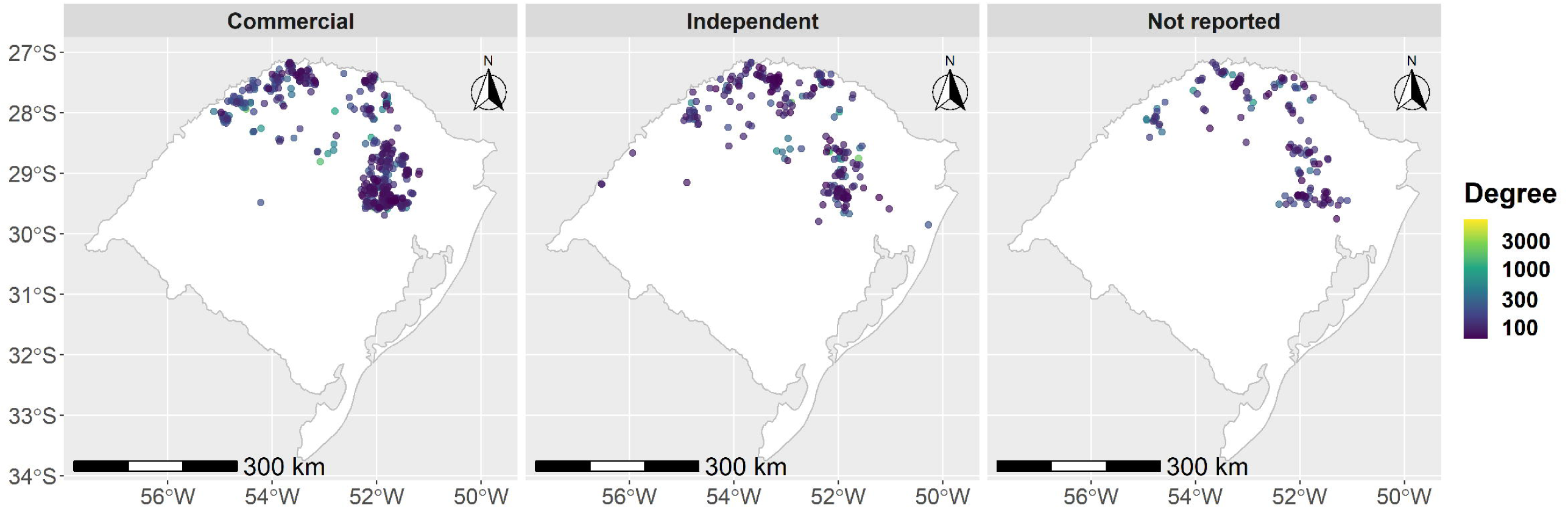
The geographic locations of the top 1,000 farms ranked by degree.

## Discussion

The majority of the movements were from commercial and independent pig farms to slaughterhouses, but we also demonstrated the presence of a contact network between independent and farms that did not report pig operation types with commercial enterprises. The later finding is of concern and a key issue in the face of the introduction and spread of foreign animal diseases (FADs) into highly connected commercial pig systems. We also demonstrated that the use of static networks to identify tightly connected farms to be selected for surveillance activities and to estimate disease spread dynamics would likely mislead disease control efforts and overestimated epidemic sizes. Thus, our model formulation accounted for the temporal order of the between-farm movements without overestimating the epidemic sizes while accounting for impact on disease spread as network-based interventions were implemented, our model also predicted the impact of network-based target disease control strategies (e.g., farm isolation, movement restriction, enhancement of biosecurity) on the accumulated number of cases over the next two years (2017 and 2018). These clearly demonstrate the advantages of temporal networks to identify key farms to compose disease control contingency plans at national, local and at production system level for vertical integrated animal production (i.e., swine, poultry), in order to quickly mitigate an introduction and subsequent spread of infectious diseases. The generation of a hot-list with farms to undergo complete lockdown or to which additional biosecurity can be rapidly applied in the face of foreign animal disease and other biological threats to national agriculture and food supply is a critical contribution of this study.

The frequency of shipments from independent to commercial pig farms reveal the importance of analyzing the interaction between pig populations of diverse biosecurity levels.

These interactions pose significant risk for the introduction and spread of infectious diseases into highly connected integrated commercial pig farms. Many studies showed that ASF spread in China and parts of the EU have been related to pig movements between non-commercial and commercial pig operations (Claire Guinat et al., 2016; Chenais et al., 2019; Miteva et al., 2020). The description of interaction between non-commercial and commercial farms remains rare in the literature (Porphyre et al., 2014). However, a study in Scotland described as much as 8.6% of movement going from small producers into commercial pig farms (Porphyre et al., 2014), meanwhile studies in North America have mostly lack the description of such movements (Kinsley et al., 2019; Sterchi et al., 2019), but for a recent a study in Iowa that described the presence of such pig flows but did not considered it their network analysis (Passafaro et al., 2019). Indeed, there is a need to capture movements between non-commercial and commercial pig farms and vice-versa, which also implies including those into the subsequent network and modeling analysis. One important limitation of this study is the fact that 21.4 % of the farms did provided information of the type of pig operation. Briefly, we evaluated the population in this category, the median number of pig was 415 (IQR 1-1441) with median number of breeding female of 72 (IQR 1-483), and median of 1 boar (IQR 0-4), see supplementary table S3 for more details. By the number of total pigs and breeding females these farms are likely smaller operation which may include farms with limited biosecurity. We highlight the need for a follow-up study to collect more information about these farms, including the type of pig operation, farm’s infrastructure and information about biosecurity. Indeed, gathering and analyzing non-commercial and commercial interactions becomes more important as the pig industry around the world prepares for the introduction of ASF, which will require tracking animal movements among all pig farms (Brown and Bevins, 2018; Jurado et al., 2018).

In this study population the static networks failed to capture time-respecting movement pathways, and consequently deceive disease control efforts. Until now a large amount of studies have considered static network views to make inferences about disease spread or to calculate outbreak sizes (Büttner et al., 2013; Lee et al., 2017; Salines et al., 2017). Mostly those studies ignored the causal fidelity of the contacts, which can be used to quantify the error of the static representation of a temporal network (Lentz et al., 2013). Thus, without such verification, it is difficult to decide if and at which time window inferences or metrics from static networks can be interpreted or used to summarize networks (Lentz et al., 2013; Büttner et al., 2016). It is dangerous to assume that aggregating contact movements into one snapshot would not overestimate or introduce edge biases (Vidondo and Voelkl, 2018). It has been suggested that to some extent, static networks provide useful information about disease related risks (Mohr et al., 2018). However, most static network studies lacked basic statistical support for their decisions for data aggregation (Büttner et al., 2013; Lee et al., 2017; Salines et al., 2017), some studies have rather mention that other studies also used similar time windows, which would support their decisions to data aggregation (Büttner et al., 2013; Lee et al., 2017; Salines et al., 2017). Finally, contact network aggregation has been a practice, even when the temporal sequences of the movements were available (Amaku et al., 2015; Cárdenas et al., 2018). Thus, the use of static networks tends to artificially connect a larger number of farms, which will consequently bias surveillance strategies to less relevant farms and compromise resource allocation, therefore when temporal movement data is available, it is strongly recommended consider such dynamic while the interest is to study the spread of diseases over time-varying networks.

The number of active pig farms was seasonal and skewed with variation of plus or minus 300 farms over the two-year period, a similar trend was observed in the volume of pigs traded. In contrast to our observations, the number of active pig farms varied largely compared with other pig populations in Germany (Büttner et al., 2016), France (Rautureau et al., 2012) and Canada (Thakur et al., 2016), even though it is difficult to compare distinct networks the comparison shown diversity especially in the volume of movements (Miele et al., 2019). Some factors that have contributed to the variations in our study may be associated with hog prices, which tends to be more volatile than in other countries. In addition, the monthly network diameter almost doubled in the middle of 2018, which seems to be a highly abnormal pattern in the network. A possible explanation for this may be related to the global configuration of the network because the nodes were connected in a way that increased the length of the shortest paths between them, which was accompanied by an increase in the GWCC. Consequently, during this period, we observed an increase in the number of nodes and connections; therefore, a given node would require more steps to reach another farm, within the network which could alone reduce the speed of disease transmission (Vidondo and Voelkl, 2018). While the GWCC increased over the years, the GSCC suffered from relevant variations throughout the months, especially in the second half of 2018, where new paths connected the GSCC with other minor components and consequently increased the size of the GWCC, which could have implications for disease transmission and persistence (Schulz et al., 2017; Marquetoux et al., 2016). The sizes of GSCC and GWCC variation were mostly attributed to variation in the number of commercial farms activities. Farms that did not report operation types represented the minority of the farms in both components, especially in April 2018 (see figure S1).

Our results show that different network-based intervention schemes successfully reduced the spread of disease independently from the local *β* transmission probability rates at the farm. We show that targeting farms by degree was the most effective intervention in reducing the number of new cases, followed by targeting farms by betweenness, which required ∼ 1,300 farms to be quarantined. Finally, as anticipated, random removal had little impact in controlling disease. A recent study explored the contribution of existing trade duration on disease transmission probabilities and the impact on epidemic sizes, suggesting that epidemic sizes were sensitive to network activity (Farine and Whitehead, 2015; Lebl et al., 2016). Indeed, we explicitly considered node activity while disease spread was simulated. Two other studies attempted to account for time-dependent disease spread while aiming to reduce *R*_0_. One study used a non-compartmental model approach and found that the number of farms that needed to be removed by degree approached 50% of all farms (Kinsley et al., 2019). The second study used a mechanistic approach in which ASF spread was simulated and found that degree ranking was the best strategy to contain the disease (Ferdousi et al., 2019). In summary, degree was shown to be the best strategy in reducing disease burden, while betweenness had the second best performance. It is worth noting that betweenness is a useful metric when studying disease diffusion (Farine and Whitehead, 2015), mainly because it is a representation of the shortest path between nodes. Here independent farms with high betweenness may bridge several commercial firms; therefore, more details about these unexplored pathways should be investigated in the future.

Based on the identified minimum number of farms of 1,000 that needed to be removed from the contact network, we simulated outbreak propagation with a deterministic SI model and identified the most efficient network-based metric capable of reducing the expected size of spread cascades. We showed that degree was the most significant intervention, with no more than 29 pig farms involved per outbreak (Figure 6). In addition, our results demonstrated that targeting more than 500 farms by degree was efficient in reducing epidemic sizes. Random surveillance was unexpectedly better than clossenes and PageRank for most of the two years of propagation. One possible explanation may be that this network is not well connected, and farms that remain to be removed may be too similar with respect to their clossennes and PageRank values; therefore, the impact on the propagation is expected to be the one resulting from random removals.

Our results have shown an uneven proportion of commercial and independent pig farms within the first 1,000 farms with the highest degree. This illustrates the interplay between the two populations, where infection can arrive at either and propagate in both directions. Here, most farms of interest lacked important biosecurity features, such as cleaning and disinfection stations (Dee et al., 2006). A study in Argentina also highlighted the lack of biosecurity on highly connected farms (Alarcón et al., 2019). On the other hand, in our study, most farms did not utilize rendering, but preferred to compost dead animals. The former was previously described as an important risk factor for the spread of porcine reproductive and respiratory syndrome (PRRS) in the US (Velasova et al., 2012; Silva et al., 2019). In the same study, composting was found to be the less risky practice when removing cull or dead pigs (Silva et al., 2019). Another relevant biosecurity point that has been described as a risk for disease introduction is the number of farm entries; here, we noticed that the majority of the farms had only one main road, which potentially represents less risk for local disease transmission (Silva et al., 2019). A disproportionate number of these highly risky farms were nurseries and breeding farms, which has often been proposed as a desired group to be targeted for surveillance (Dorjee et al., 2013). In addition to the identification of priority farms, the identification of relevant biosecurity gaps at those farms is invaluable, especially under the current preparedness for the introduction of foreign animal diseases such as ASF. One of the best options is the implementation of enhanced biosecurity plans, such the Secure Pork Supply (SPS) Plan, designed to provide business continuity in the event of a foreign animal disease outbreak as well as help protect operations from endemic diseases (Pudenz et al., 2019).

Finally, more comprehensive studies coupling animal movements with local and environmental routes of transmission while accounting for farms biosecurity and infrastructure are needed to further explore the mechanisms of pig disease propagation, for which a complete multiscale mechanistic model is necessary (Qi et al., 2019).

### Considerations

The use of static networks to target key farms to control the disease from spreading should be interpreted with caution due to several limitations associated with animal movement aggregation, as shown in this study and elsewhere (Lentz et al., 2016). An obvious issue here is to assume that the same pair of farms would trade with each other throughout the entire study periods.

Therefore, we argue that the causal fidelity of the temporal network should be presented before considering the use of the static networks in the development of disease surveillance or control strategies (Lentz et al., 2016; Payen et al., 2019). Most of recent studies did not account for this issue (Büttner et al., 2013, 2018; Marquetoux et al., 2016; Motta et al., 2017; Alarcón et al., 2019), or considered the temporal network pathway accessibility of each farm.

In this study, we did not examine feed-truck or other vehicle movements or sharing. Such information is absent from the majority of the current literature (Lentz et al., 2016), but it has been included in some studies (Augusta et al., 2019; Sterchi et al., 2019). It is critical to acknowledge that the movement of feed trucks and short-distance transportation of breeding replacement animals or culled sows contributed to disease transmission (Silva et al., 2019). Indeed, before these additional contacts are not taken into consideration, our understanding of disease spread pathways will likely continue to be limited.

Finally, the modeling framework proposed here can be broadly used to calculate the minimum number of farms to be targeted for the control of disease spread when temporal contact data are available and to determine the network metrics that should be preferred for selection of target farms according to the approximation of epidemic sizes. In addition, the proposed framework could be used as a decision tool for both local official veterinary services and the private sector for evaluating animal movements and developing a more risk-oriented surveillance strategies. At production system level, it allows for the creation of a hot-list with farms to which the reinforcement of biosecurity would be key in the face of an outbreak. The model also provides an estimation of the amount of personnel and material resources that would need to be allocated to control the epidemic, which is key for preparedness for future events (Casal et al., 2019). Future studies are also needed to explore the contribution of movements to slaughter houses, especially when pig are redirected to other slaughter houses (i.e., slaughter not able to slaughter all animals) or send back to the farm of origin, this is very common in the US, however little is known relevance in spreading disease back to the farm, however there is some early discussion about it (Russell et al., 2020).

## Conclusion

The contact among farms of distinct biosecurity by pig movements identify a potential gap that could result in the introduction and dissemination of FAD into commercial and highly connected pig populations. One-third of all pig farms traded with the same partners in the course of two years; consequently, the static networks did not offer reasonable information about the realized temporal movement pathways. In this scenario, the use of static networks to target vulnerable or high-risk farms by network metrics would likely overestimate the role of those farms in disease transmission and limit the accuracy needed to optimize surveillance. Our proposed modeling framework provides a parsimonious solution to this problem; it handles the temporal order of movements, calculates the minimum number of farms needed to be targeted and approximates the expected epidemic size in a worst-case scenario where each movement can effectively transmit disease. The results from the application of this approach identified priority pig farms that had remarkably limited biosecurity. This fact imposes important vulnerability to pathogen introduction and spread; therefore, those farms could benefit from the implementation of enhanced biosecurity plans, such as what has been promoted and implemented in the US by SPS plan. Finally, the introduction of risk-based approaches may optimize the costs of routine surveillance and can be used in preparation for the introduction of FAD into the local pig industry.

## Acknowledgements

This work was supported by the Fundo de Desenvolvimento e Defesa Sanitária Animal (FUNDESA-RS) and by the NC State University, College of Veterinary Medicine, Global Health funds.

## Authors’ contributions

GM, FPNL, JV and AARM coordinated movement data collection. GM conceived paper ideas. GM, JAG and NCC participated in the design of the study. NCC and GM conducted data processing and cleaning. GM, JAG and NCC wrote and modified computer algorithms and conducted analysis. GM, JAG and NCC developed simulation models. GM and NCC wrote the manuscript and JA edited the manuscript. All authors discussed results and commented on the manuscript.

## Ethical statement

The authors confirm that the ethical policies of the journal, as noted on the journal’s author guidelines page. Since this work did not involve animal sampling neither questioner data collection there was no need for ethics permits.

## Data Availability Statement

The data that support the findings of this study are available from the regional veterinary office. Restrictions apply to the availability of these data, which were used under confidentiality agreements.

