## Supplementary material for "Quantifying the dynamics of pig movements improves targeted disease surveillance and control plans": ss

**Running title: Time-varying networks improve surveillance in pigs**

Gustavo Machado[1*], Jason A. Galvis[1], Francisco P.N. Lopes [2], Joana Voges[2], Antônio A.R. Medeiros [2], Nicolas C. Cárdenas [3]

[1]Department of Population Health and Pathobiology, College of Veterinary Medicine, Raleigh, North Carolina.

[2] Secretary of Agriculture, Livestock and Agribusiness of State of Rio Grande do Sul (SEAPI-RS), Porto Alegre, Brazil.

[3]Department of Preventive Veterinary Medicine and Animal Health, School of Veterinary Medicine and Animal Science, University of São Paulo, São Paulo, Brazil.

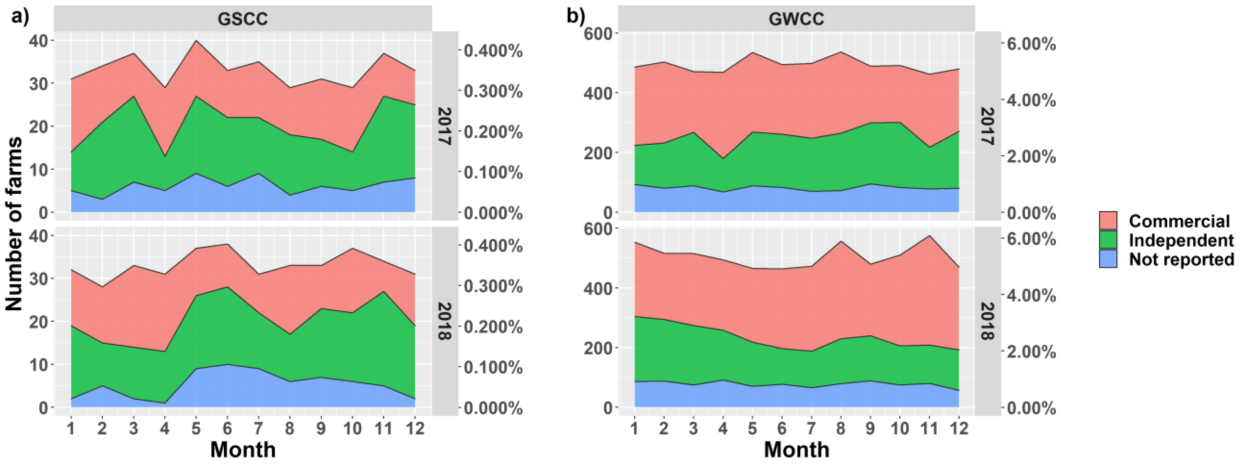


**Figure S1.** The monthly GWCC and GSCC for each farm operation type, commercial, independent, and not reported.

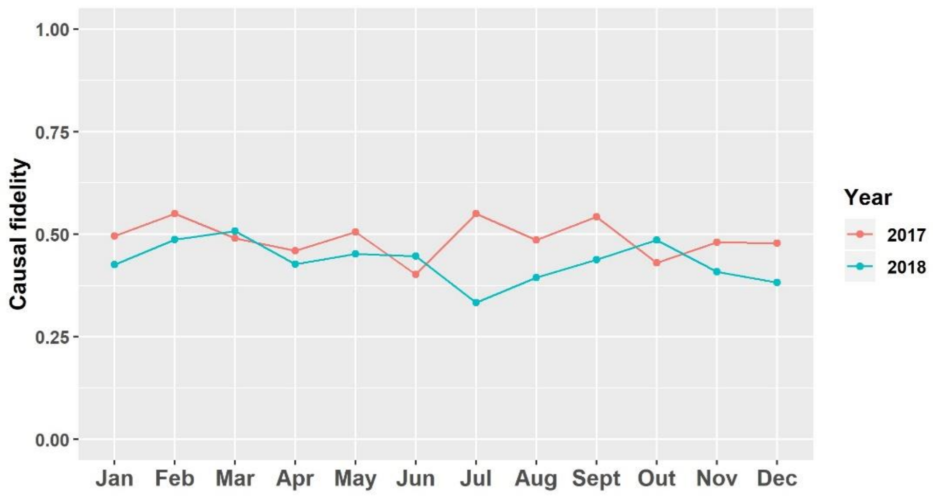


**Figure S2.** Monthly causal fidelity.

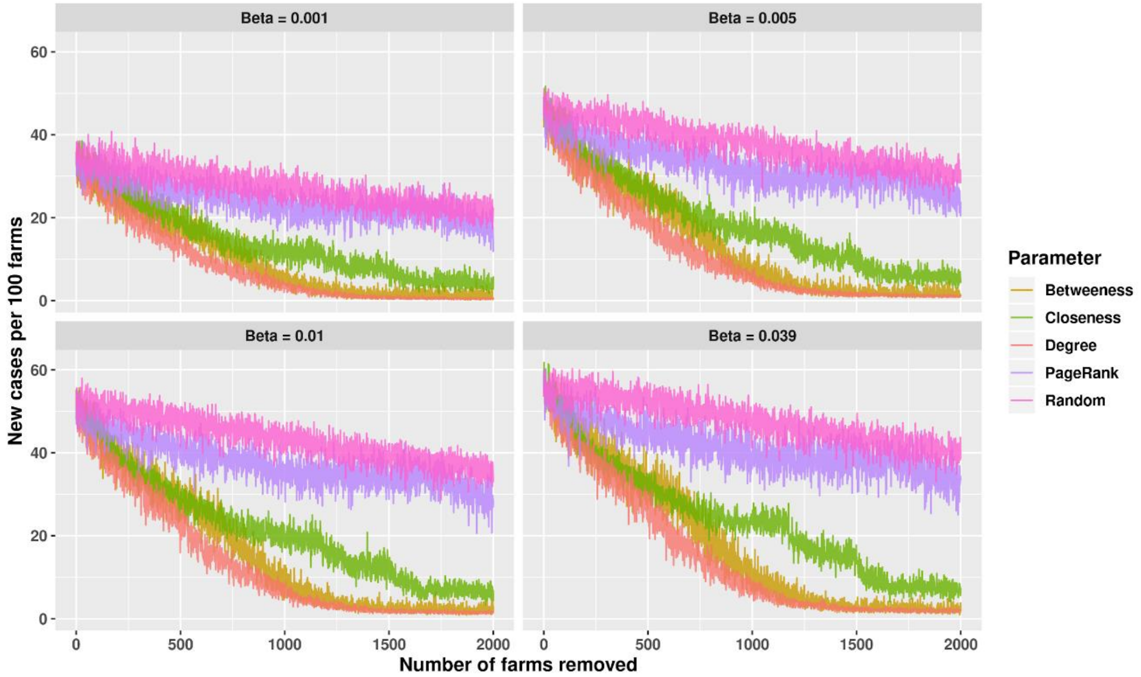


**Figure S3**. Simulated disease spread dynamics within the swine contact network. Simulations assumed 950 pig farms (10% prevalence) were infected at day "0". The simulations assumed a farm prevalence of 0.1%.

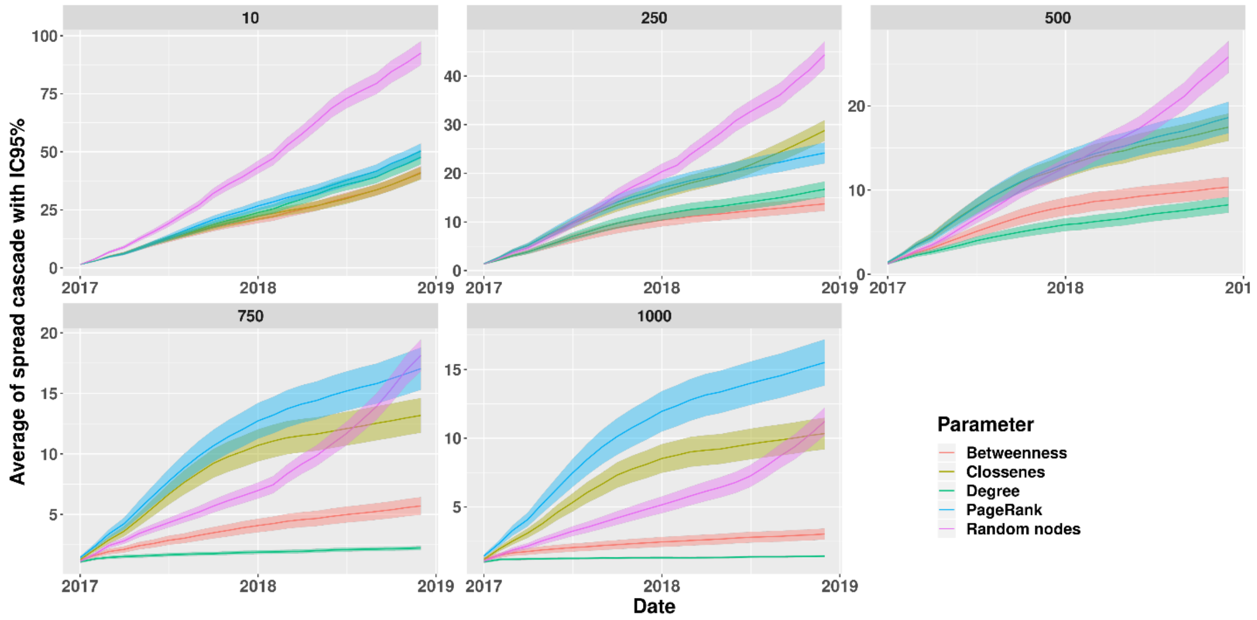


**Figure S4**. Epidemic size by node removal. The y-axis shows the cumulative size of the spread cascades from January 2017 through December 2018 for five scenarios. On day "0", 10, 250, 500, 750 or 1,000 farms were removed from the simulation, while the temporal network metrics were calculated for all remaining farms. The lines is the average values and shaded ± 95% CIs.

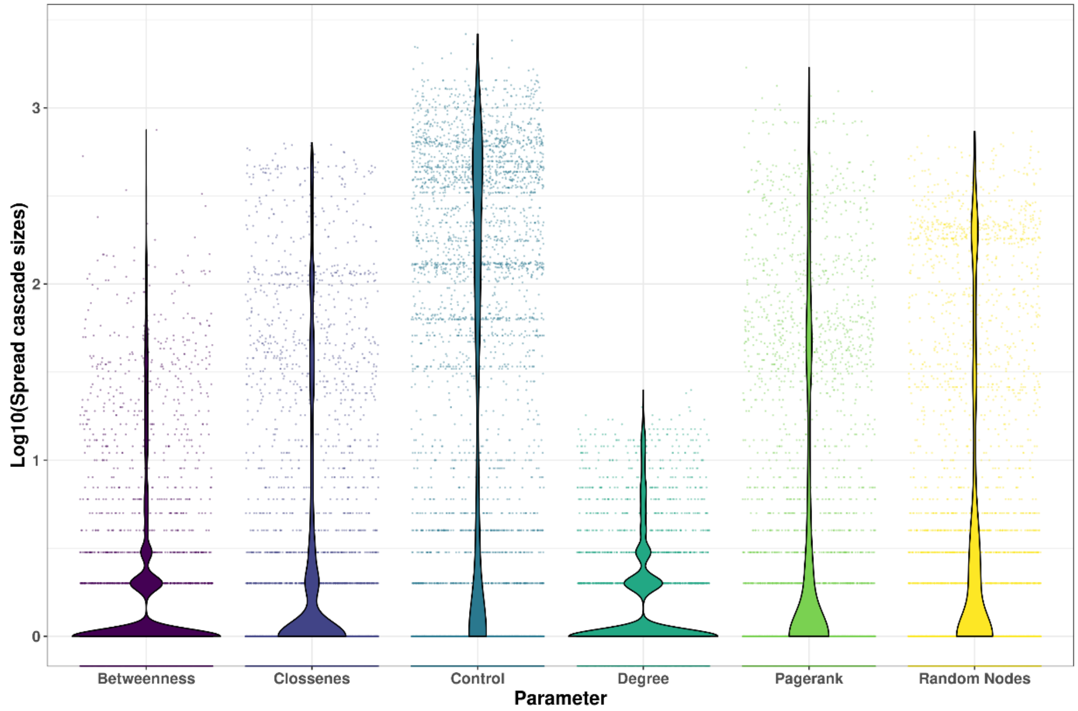


**Figure S5**. The accumulated spread cascades size by intervention metrics. Violin plot areas represent kernel densities distribution and dots log transformed spread cascade.

**Table S1**. Movement between commercial and non-commercial farms. Commercial are further divided into commercial those farms with contract with integrated pig companies, and independent are commercial but has not active contract with integrated pig companies, but also include small farm holder that can be considered backyard producers. Not reported are farms that fail to report the local authorities their farm business type. The summary show number of pigs and number of movements from 2017 to 2018 are shown.

| **Farm of origin** | **Farm of destination** | **Number of animals** | **Number of shipments** | **Proportion of total movements** |
|---|---|---|---|---|
| Commercial | Slaughterhouse | 9983618 | 121598 | 34.60 % |
| Independent | Slaughterhouse | 4675247 | 65971 | 18.77 % |
| Commercial | Commercial | 13936172 | 51495 | 14.65 % |
| Independent | Independent | 5757826 | 30888 | 8.79 % |
| Not reported | Slaughterhouse | 1605273 | 27206 | 7.74 % |
| Commercial | Independent | 5131285 | 21566 | 6.14 % |
| Not reported | Independent | 2504556 | 14036 | 3.99 % |
| Independent | Commercial | 2990437 | 11774 | 3.35 % |
| Not reported | Commercial | 1933443 | 6939 | 1.97 % |

**Table S2** The size of GSCC and GWCC for the full two years static network, in which the proportion of farm types is considered. We show component sizes by pig production type and for pig operation classification.

| **Farm type** | **GSCC size** | **GWCC size** | **GSCC %** | **GWCC %** |
|---|---|---|---|---|
| Commercial | 260 | 3157 | 2.75 | 33.42 |
| Independent | 401 | 2670 | 4.25 | 28.27 |
| Not reported | 202 | 1732 | 2.14 | 18.34 |

**Table S3** The number of pig by type of operation

| **Pig operation type** | **Total number of pigs** | **Number of breeding female** | **Number of boars** |
|---|---|---|---|

| | | | |
|---|---|---|---|
| Commercial | Median 2,087 (IQR 933-6,707) | Median 800 (IQR 285-1,890) | Median 5 (IQR 2-9) |
| Independent | Median 949 (IQR 220-2,866) | Median 192 (IQR 25-678) | Median 1 (IQR 0-5) |
| Not reported | Median 415 (IQR 62-1,441) | Median 72 (IQR 6-483) | Median 1 (IQR 0-4) |

**Table S4-** Comparison between epidemic sizes at the end of two years spread when 1.000 farms are removed from the network.

| **Pair comparison** | **Z-score** | **P-unadjusted value** | **P-adjusted value** |
|---|---|---|---|
| Degree - Pagerank | 217,320,480,005,551 | 1.02E-90 | 1.53E-89 |
| Degree - Clossenes | 197,067,056,182,027 | 1.89E-72 | 1.42E-71 |
| Degree - Betweenness | 187,547,323,255,671 | 1.77E-64 | 8.86E-64 |
| Control - Pagerank | 165,600,224,174,183 | 1.36E-47 | 5.08E-48 |
| Degree - Random Nodes | 14,349,662,207,118 | 1.07E-32 | 3.21E-32 |
| Clossenes – No control | -14,201,488,085,431 | 8.97E-32 | 2.24E-31 |
| Betweenness – No control | -131,354,909,052,691 | 2.06E-25 | 4.42E-25 |

| | | | |
|---|---|---|---|
| Pagerank - Random Nodes | -820,592,758,572,372 | 2.29E-02 | 4.29E-04 |
| Degree – No control | 804,579,970,156,665 | 8.57E-02 | 1.43E-01 |
| No control - Random Nodes | 78,082,826,340,933 | 5.80E-01 | 8.70E-01 |
| Clossenes - Random Nodes | -618,271,575,570,572 | 6.30E+04 | 8.59E+03 |
| Betweenness - Random Nodes | -534,440,148,004,969 | 9.07E+06 | 1.13E+07 |
| Betweenness - Pagerank | 252,132,352,034,643 | 0.0116914305133186 | 0.0134901121307522 |
| Clossenes - Pagerank | 184,417,635,777,645 | 0.0651574427029602 | 0.0698115457531717 |
| Betweenness - Clossenes | 0.700670193306622 | 0.483508861646078 | 0.483508861646078 |
